# Rethinking De-Extinction Criticism: A Multi-Dimensional Model for Prioritizing Revivable Species under Funding Controversies

**DOI:** 10.1101/2025.07.16.665200

**Authors:** Alper Kaan Selçukoğlu, Deha Kaykı

## Abstract

De-extinction is often criticised for siphoning funds from classical conservation, yet quantitative evidence remains scarce. Here we assemble a disaggregated United States finance dataset covering 2021 to 2024 and find that every dollar invested in de-extinction originated from private sources and coincided with a net rise in public and NGO conservation budgets. Rather than displacing existing capital, de-extinction mobilised new money and produced a crowding-in effect that enlarged the pool available for biodiversity protection.

To channel these additional resources strategically, we develop the Multidimensional De-extinction Effectiveness Model (MDEM), which ranks candidate species by ecological benefit, genetic insight, technical feasibility, and habitat readiness. Applied to sixteen taxa, MDEM consistently places the thylacine (*Thylacinus cynocephalus*) first; Monte Carlo tests (10,000 runs, *±* 3 input jitter) yield a median class-level stability of 73.2%, underscoring robustness to uncertainty.

Our results position de-extinction as a complementary branch of conservation finance and offer a transparent framework for steering biotechnology investments toward species with the greatest combined ecological and scientific return.

## 1 Introduction

The accelerating biodiversity crisis compels conservation science to reassess both its methodological foundations and strategic toolkit. While conventional conservation has been effective in protecting extant species and habitats in specific contexts, it has largely focused on diagnosing ecological problems rather than developing scalable, future-oriented solutions [1]. Consequently, the long-term ecological and financial sustainability of traditional conservation models is under increasing scrutiny, driving calls for innovation and systemic reform.

Against this backdrop, de-extinction has emerged as one of the most provocative proposals on the table. The term describes the use of modern biotechnology to restore ecological functions and phenotypic traits once performed by extinct species, without insisting on perfect genetic replication [4, 6]. The concept itself is not new; its origins trace to the early 20th century, most notably the Heck brothers’ attempt to recreate the extinct aurochs through selective breeding [3]. Although both theoretical and practical efforts had emerged before, the 2021 founding of Colossal Biosciences, a U.S.-based company, marked a key shift by turning the concept into a globally visible biotech initiative [5]. Today, these efforts stand at the center of an intense and polarized debate. Proponents argue that de-extinction can stimulate public interest, drive technological innovation, and broaden both the conceptual and practical boundaries of conservation science [11]. Critics, by contrast, raise ethical concerns and caution that such projects might divert scarce resources from the protection of extant species. They emphasize potential animal welfare issues related to cloning and surrogacy, the unpredictable ecological consequences of introducing proxy species into existing ecosystems, and the risk of unforeseen disruptions to current ecological balances [12, 10, 9]. Furthermore, they warn that de-extinction initiatives may siphon funding, attention, and institutional support away from conservation efforts targeting currently endangered species.This concern is frequently framed as the “crowding-out hypothesis,” which assumes that conservation budgets are fixed, opportunity costs narrowly defined, and indirect benefits such as public engagement or technological advances of minimal relevance.

Beyond academic discourse, public and media narratives have played a significant role in shaping perceptions of de-extinction. As its profile in mainstream media has expanded, de-extinction has at times been incorrectly portrayed as a rival to traditional conservation efforts. This binary framing perpetuates the misleading notion that these approaches are inherently in conflict, rather than recognizing their potential complementarity within an integrated strategy to protect biodiversity.

Despite extensive media portrayals, significant knowledge gaps persist. While research on de-extinction has evolved beyond its initial ethical debates, the field still lacks clear frameworks to determine which projects should be prioritized. Most financial assessments to date have treated conservation holistically, seldom distinguishing between public and private funding streams. However, because de-extinction efforts are almost entirely supported by private capital, the critical question is not whether they displace public conservation budgets, but whether these funds are being directed toward ecologically and scientifically meaningful species. Existing cost-effectiveness models primarily emphasize ecological benefit relative to cost, frequently neglecting essential dimensions such as genetic knowledge, technical feasibility, and habitat readiness. In the absence of improved frameworks, it remains unclear how to identify and prioritize the most valuable de-extinction initiatives.

De-extinction projects are capital-intensive, technically complex, and highly visible, making it essential to close these knowledge gaps as more than a purely theoretical exercise. Advancing responsibly requires robust methods to assess both the ecological benefits and the scientific and technological value of candidate projects. This study addresses a central question: how can project selection be improved so that private funding is directed toward the most meaningful and impactful cases?

To address these issues, we analyzed conservation funding data from 2021 to 2024, comparing investments across three domains: classical conservation, conservation biotechnology, and de-extinction. Our findings suggest that de-extinction funding expands rather than diminishes overall conservation resources, challenging the crowding-out hypothesis and instead pointing to a crowding-in dynamic. These results underscore the need for more rigorous tools to guide the allocation of this expanded pool of resources.

In response, we introduce the Multidimensional De-Extinction Effectiveness Model (MDEM), a prioritization framework that integrates ecological benefit, genetic potential, technical complexity, and habitat suitability. This approach links macro-level funding trends with project-level decision-making, helping ensure that new investments are directed toward the most ecologically and scientifically meaningful cases.

Building on prior debates—including the influential analysis by [2]-we contend that several core assumptions underlying critiques of de-extinction are rarely valid in practice. These include the notions that conservation budgets are rigidly fixed, that resurrected species would require identical management to extant species, and that indirect benefits such as public engagement and technological innovation are negligible. We demonstrate that adopting a broader, multidimensional framework provides a more robust foundation for setting priorities within both de-extinction and conservation biotechnology.

## 2 Background and Conceptual Framework

Conservation projects have traditionally been ranked by their expected ecological gain per unit of financial investment. While this remains a key metric, recent evaluations of Europe’s LIFE-Nature funds and broader species prioritization literature demonstrate that reliance on ecological criteria alone is insufficient [8, 7].

A key challenge is that conservation decisions often fail to incorporate factors such as a species’ evolutionary and ecological uniqueness, its functional importance within ecosystems, its symbolic or cultural significance, the cost of management, and the likelihood of success. Neglecting these considerations can result in inefficient use of scarce conservation resources and biodiversity losses that might otherwise have been prevented. At the same time, a significant portion of conservation funding is often directed toward so-called vanity species, which are charismatic yet non-threatened animals that capture public and donor attention but contribute little to solving the broader biodiversity crisis [18]. This bias is further supported by recent studies, which indicate that although only approximately six percent of officially threatened species have received conservation funding, nearly twenty-nine percent of total funds have been directed to species classified as of least concern [13].

Importantly, similar mismatches are observed not only in species-level prioritisation but also in habitat and area-based conservation efforts, where an exclusive focus on expanding protected areas has often failed to address deeper ecological and management challenges. Assessments conducted between 2010 and 2019 underscore these persistent issues. Coverage remains far from ecologically representative: nearly 80 percent of approximately 12,000 threatened species lack adequate protection, and more than 1,400 species receive no protection at all [14]. Management effectiveness is frequently low, driven largely by chronic underfunding and weak enforcement, while integration with Indigenous and local conservation initiatives remains limited. These shortcomings indicate that merely increasing the extent of protected areas will not suffice to halt biodiversity loss or safeguard essential ecosystem services.

These limitations demonstrate that expanding protected areas alone is insufficient to meet contemporary biodiversity goals. Additional strategies are needed, particularly those leveraging novel tools from conservation biotechnology to restore species and their ecological functions. Conservation biotechnology encompasses methods such as gene editing, synthetic biology, assisted reproduction, and advanced veterinary interventions designed to support species recovery, rebuild populations, and enhance ecosystem resilience [15, 17]. Early evidence suggests that these emerging techniques can help rescue species that conventional conservation has been unable to protect, functioning in complement to rather than as a replacement for established approaches.

One example from conservation biotechnology is the use of assisted reproductive technologies (ART), such as artificial insemination and in vitro fertilization, in conservation breeding [19, 16]. These methods aim to improve the genetic health of endangered populations. While ART has led to births in over fifty species, its use in long-term population recovery is still rare, with only a few well-known successes like the giant panda and black-footed ferret [19]. This shows both the promise and current limits of such tools, especially when used alongside other conservation strategies.

As part of this broader picture, modern de-extinction can be seen as part of conservation biotechnology. It can be classified as a branch of this broader field, yet it also has a distinct focus, as it aims to bring back extinct species while contributing to wider conservation and ecosystem restoration goals. This dual focus gives de-extinction a unique place within conservation science, combining scientific innovation with ecological and symbolic impact. De-extinction projects aim to help rebuild lost ecological functions, advance genomic and reproductive technologies, and inspire public interest and funding. When seen this way, de-extinction complements other biotechnological approaches and helps strengthen overall conservation efforts.

However, despite these promises, de-extinction has also faced important criticism. Authors like [2] and [11] have expressed concerns that such projects could divert limited conservation resources away from existing species and might ultimately lead to further biodiversity loss. Critics also highlight broader risks, such as creating a false sense that technology can solve conservation problems, reducing public motivation to protect endangered species, and introducing ecological uncertainties we do not fully understand. At the center of these debates are key questions about whether conservation budgets are fixed, whether funding can easily move between different types of projects, and whether revived species would need the same management as living ones. Exploring these points is essential for understanding what role de-extinction can realistically play in future biodiversity efforts.

## 3 Methods

To empirically assess whether de-extinction initiatives crowd out or complement conservation efforts, we conducted a two-part analysis combining quantitative funding data with a multidimensional prioritization framework. First, we compiled and analyzed disaggregated conservation funding records from 2021 to 2024, focusing on the United States and major global organizations such as the IUCN and WWF, and distinguishing among classical conservation, conservation biotechnology, and de-extinction-focused initiatives. Second, we developed and applied an extended effectiveness model that builds upon but diverges from classical cost-effectiveness approaches such as that of [2], integrating additional ecological, scientific, and symbolic dimensions. Prior to classification, we manually assembled a bespoke database of financial records from publicly available reports, institutional disclosures, and corporate statements for the period 2021–2024; the complete dataset is provided in Supplementary Table 1, ensuring full traceability of all entries included in the analysis. Graphical visualizations, including stacked bar charts and pie charts, were prepared using Python’s Matplotlib library. Where appropriate, SciPy and NumPy were used for curve fitting and graphical adjustments. All visualizations were designed to maintain consistency, clarity, and interpretability across sectors and years

### 3.1 Funding Source Classification

To compare funding streams systematically, we categorized each financial record by funding source, using publicly available information. We defined four main categories: private venture capital, foundation or NGO grants, public (governmental or multilateral) funds, and mixed or hybrid philanthropic models. If an investor’s background could not be confirmed, the entry was labeled as ‘Uncertain’ and excluded from the source-type analysis. This classification helped us assess the origins and diversity of financial inputs and compare the funding patterns of traditional and synthetic approaches to biodiversity conservation.

### 3.2 Analytical Tier Definitions

For the purposes of this study, we treated de-extinction as analytically distinct from the broader field of conservation biotechnology, even though it is technically a subfield. This distinction is justified by the explicit self-identification of companies like Colossal as ‘de-extinction companies’ and by the unique public and scientific debates surrounding species resurrection. Accordingly, funding records were grouped into three analytical categories: Tier 1, de-extinction initiatives (for example, Colossal); Tier 2, conservation biotechnology projects (such as Revive & Restore and BioRescue); and Tier 3, classical conservation efforts (including grants from the US Fish and Wildlife Service, WWF, IUCN, and USAID).

### 3.3 Scope Definition and Assignment

We introduced a custom variable, ‘Scope,’ to capture the geographic and operational reach of each funding stream, defined as Domestic (local or sub-national grants), National (nationwide programs), Multinational (formal cooperation between countries), or Global (programs with stated or implied worldwide coverage). Classifications were assigned based on the mandates and publicly stated objectives of implementing organizations. While ‘Scope’ was recorded for all entries, it was not used in the main analysis and is provided for reference in Supplementary Material.

### 3.4 Inclusion and Exclusion

To maintain analytical consistency, we included only funds explicitly designated for terrestrial biodiversity conservation. Included were species-specific or multi-species terrestrial programs and national or subnational recovery initiatives for threatened species. We note that de-extinction projects, despite involving complex technological and ecological dimensions, are typically framed and funded as species-focused initiatives rather than as general habitat or ecosystem programs. This focus aligns them analytically with classical species conservation efforts and justifies their inclusion in the terrestrial species conservation category. Excluded from the analysis were marine-only conservation funds, general environmental or ecosystem restoration projects not directly linked to species conservation, habitat protection projects without a species focus, and aggregate budget lines already broken down in annual calls to avoid double counting.

### 3.5 Handling Multi-Year and Aggregated Data

Multi-year program envelopes were excluded when sub-grants or annual calls were already accounted for. When multi-year programs covered both biodiversity and non-biodiversity components, only the portion explicitly designated for species conservation was included. Where disaggregation was impossible, the entry was omitted.

To standardize temporal analysis, multi-year funds were annualized using the formula:

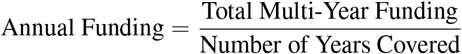

For example, a program distributing $50M between 2022 and 2025 would contribute $12.5M per year. While the primary analytical window covered the years 2021–2024, we included the proportional share of multi-year programs extending into 2025 to ensure that ongoing funding streams were adequately represented.

### 3.6 Currency Conversion and Regional Notes

While the overall aim of this study was to assess conservation funding dynamics at a global scale, practical constraints limited the scope of data extraction. Although substantial public conservation investments exist in regions such as Europe, Australia, China, and Brazil, many national and regional funds are not regularly disaggregated or available in a form suitable for cross-comparison. In particular, many public records combine area-based and species-focused budgets without clear separation. To ensure analytical consistency and generate a clean dataset, we focused on funding streams with reliable, publicly available, and regularly updated records, primarily from the United States as well as international organizations such as the IUCN and WWF. The analysis period began in 2021, corresponding to the founding of Colossal, which marked a turning point in de-extinction investments. Data from the year 2025 were excluded due to the absence of standardized classical conservation funding reports for that year. This choice necessarily excluded some significant national funding sources but ensured the comparability, traceability, and robustness of the analysis.

### 3.7 Funding Estimation Procedure

To estimate USAID’s FY2024 biodiversity-specific funding, we relied on publicly available congressional budget summaries and environmental spending reports. Although the FY2024 omnibus appropriations bill allocates a total of $1,050,000,000 in bilateral funding across four programmatic areas—biodiversity, sustainable landscapes, climate adaptation, and clean energy—it does not disaggregate the allocation by individual category.

Given that in FY2023, USAID’s biodiversity portfolio was funded at approximately $375,000,000, we inferred its likely FY2024 allocation by calculating its proportional share within the previous year’s total. Assuming that the combined budget for the four areas in FY2023 was $1,105,000,000 (i.e., FY2024’s $1,050,000,000 plus the reported $55,000,000 reduction), the biodiversity share constituted approximately 33.9% of that total. Applying this proportion to the FY2024 combined allocation yields an estimated $356,000,000 for biodiversity-related programs.

This method assumes proportional continuity in program-level prioritization from FY2023 to FY2024, which we consider a reasonable approximation in the absence of contradictory evidence or program-specific appropriations. However, due to the lack of disaggregated figures in the official FY2024 documentation, this estimate should be interpreted as an approximate proxy rather than an exact value.

### 3.8 Methods: Multidimensional De-Extinction Model

This section (§3.8) details the modelling workflow used to rank candidate species based on multiple ecological, scientific, and economic dimensions. The following subsections describe the key steps and criteria applied in the prioritization model.

### 3.9 Variable Normalisation

To ensure commensurability, every raw input *X* (scientific feasibility, cost, *etc*.) is mapped onto [0, 1] by min–max scaling:

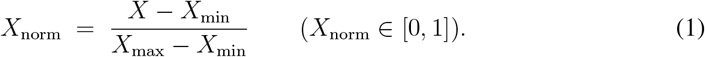

### 3.10 Equal-Weight Aggregation

Given *n* dimensions per species (*X*_1…*n*_), equal weights are assumed in the absence of expert priors:

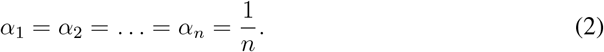

The composite (unscaled) priority score becomes

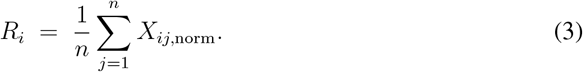

### 3.11 Composite Cost Modelling

Total cost combines laboratory complexity and habitat compatibility.

#### 3.3.1 Technical baseline

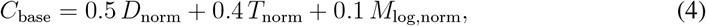

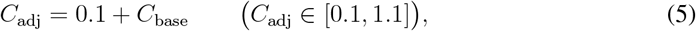

where *D* = genetic divergence, *T* = time since extinction, *M* = body mass (log-scaled). A floor *ε ∈* [0.05, 0.10] avoids division by near-zero values. The weighting scheme in Eq. (4) assigns 0.5 to genetic divergence (*D*), 0.4 to time since extinction (*T*), and 0.1 to log-scaled body mass (*M*_log_). Genetic divergence carries the highest weight, reflecting its central role in determining technical feasibility, as higher genomic dissimilarity increases challenges in nuclear-mitochondrial compatibility, epigenetic reprogramming, and developmental viability. Time since extinction is weighted slightly lower but remains critical due to habitat shifts, ecological vacancy, and societal familiarity. Body mass, log-transformed to reduce skew, is given a minor weight to acknowledge the practical complications posed by large-bodied species without letting this factor dominate the score. The adjustment term *ϵ ∈* [0.05, 0.10] ensures numerical stability by preventing near-zero denominators.

#### 3.3.2. Habitat compatibility

To account for ecological feasibility, we implemented a habitat mismatch risk factor:

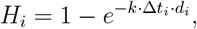

where Δ*t*_*i*_ denotes time since extinction and *d*_*i*_ is the species-specific habitat degradation score ([0, 1]), derived from land-use databases or expert consensus. Parameter *k* was calibrated so that the species with the highest Δ*t*_*i*_ *· d*_*i*_ combination reached *H*_target_ = 0.95. This ensures that species with long extinction durations but intact ecosystems (e.g., Arctic or boreal systems) are not over-penalized, while recently extinct species with irreversibly degraded habitats are downgraded accordingly.

Habitat degradation scores (*d*_*i*_) were assigned on a 0 to 1 scale, where 0 indicates no significant habitat loss or degradation, and 1 indicates near-total loss or severe alteration of the original ecosystem. Scores were determined by the authors based primarily on historical land cover transformation (such as deforestation, agricultural expansion, or urbanization) and current habitat integrity, including fragmentation, invasive species pressure, pollution, or human settlement density. For example, tundra-steppe regions were scored at 0.1, reflecting minimal transformation since the Late Pleistocene; areas like Tasmania’s forests were scored at 0.3, reflecting moderate land conversion, increasing human population, and invasive species pressure; intermediate scores like 0.6 were assigned to regions such as the former habitat of Neovison macrodon, reflecting more substantial human impact; and tropical islands like Mauritius were scored at 0.9, reflecting severe alteration from tourism, invasive species, and dense human activity. Full species-specific scoring details are provided in Supplementary Table S1

### 3.12 Multidimensional Effectiveness Score

The final score for species *i* is

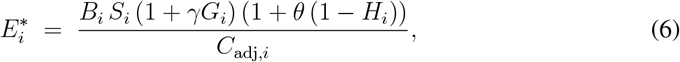

where *B* = biodiversity value, *S* = symbolic/strategic value, *G* = scientific insight, *γ* = science weight, *θ* (default 0.2) penalises habitat risk.

### 3.13 Priority Classification

Species are assigned to priority bands via quartiles of the 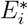 distribution: *High* (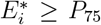), *Medium* (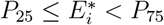), and *Low* (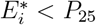).

### 3.14 Illustrative Comparison and Model Interpretation

To demonstrate the functioning and interpretability of the Multi-Dimensional Effectiveness Model (MDEM), we conducted an illustrative comparison using two well-known candidate species: *Thylacinus cynocephalus* (thylacine) and *Mammuthus primigenius* (woolly mammoth).

#### Scoring Framework

Scoring Framework. Biodiversity value (Bi) was assigned based on five semi-quantitative ecological criteria: (1) endemism, (2) functional uniqueness, (3) phylogenetic distinctiveness, (4) ecosystem engineering capacity, and (5) ecological vacancy. While standardized indices (e.g., EDGE or FUSE scores) were not directly applied due to data limitations, author-assigned scores were derived from structured literature review (see Supplementary Table S1).

Strategic or symbolic value (Si) was evaluated using metrics such as media coverage, cultural icon status, and public familiarity. Scientific or technological contribution (*G*_*i*_) reflected the potential of the species to advance genomic tools, cellular models, or functional genomic understanding. This was assessed based on factors such as CRISPR applicability, genome assembly status, and the extent of existing biotechnological or biological research on the species itself or closely related taxa.

To account for subjectivity in author-assigned scores, we conducted Monte Carlo simulations to assess the stability of species prioritisation under input uncertainty. Specifically, we applied uniform jitter of ±3 points across 10,000 replicates to three semi-subjective variables: biodiversity value (Bi), strategic or symbolic value (Si), and scientific or technological contribution (Gi). These parameters were selected for perturbation because they involved partial author judgement during initial scoring, unlike variables such as habitat adjustment (Hi) and adjusted cost factor (Cbase adj), which were based on quantitative sources and held fixed. Input scores were scaled on a 1–10 range, making the ±3 jitter equivalent to roughly 30 percent of the total range and providing a deliberately conservative estimate of plausible variation. Additional tests with lower jitter amplitudes (±1, ±2) showed negligible effects on rank stability, confirming the robustness of the ±3 choice. This selective perturbation approach allowed us to estimate how sensitive final priority rankings were to subjective inputs, without artificially inflating total model variance. Results are summarized in Figure 4 and detailed in the Results section

#### Comparative Input Values

| Species | $B$ | $S$ | $G$ | $\Delta t$ (yrs) | $D$ (My) | $M$ (kg) | $d_i$ |
| --- | --- | --- | --- | --- | --- | --- | --- |
| Thylacine | 10 | 9 | 10 | 90 | 38.2 | 17 | 0.03 |
| Mammoth | 9 | 9 | 10 | 4,000 | 6.0 | 6,400 | 0.10 |

Here, the thylacine receives a high biodiversity (*B*) score, primarily reflecting its pronounced phylogenetic distinctiveness, with an estimated divergence time of 38.2 million years from its closest living relatives, as well as its crucial ecological role as an apex predator in the Tasmanian ecosystem prior to extinction. Importantly, the thylacine also holds significant conservation relevance, as insights gained from its recovery efforts could inform broader marsupial conservation strategies and enhance our understanding of stress physiology, reproductive biology, and adaptive genetics within this highly imperiled mammalian lineage.

The mammoth, in contrast, scores slightly lower in *B* due to a comparatively shallower divergence time of approximately 6.0 million years, yet it attains an equally high score in scientific or technological contribution (*G*), reflecting not only its iconic position within paleogenomics and de-extinction research but also its potential to inform and advance conservation efforts for extant elephantid species and other large mammals under threat. This comparison illustrates how the framework integrates both ecological and technological dimensions, ensuring that candidates are not solely ranked by biological distinctiveness but also by their capacity to contribute to contemporary conservation science..

#### Model Outcomes

Baseline scores (without habitat adjustment):

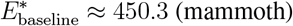

With habitat adjustment (*θ* = 0.2):

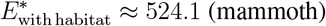

This yields a relative score increase of approximately 16.4% for the mammoth, highlighting that species with relatively intact or recoverable habitats benefit meaningfully from the compatibility adjustment. The adjustment amplifies the ecological feasibility signal, ensuring that candidates are not penalized solely for technical or symbolic dimensions, but are evaluated within a realistic habitat restoration context.

The habitat adjustment factor thus allows the model to meaningfully distinguish between species based not only on historical significance or technical feasibility, but also on realistic ecological reintroduction potential. Importantly, the mammoth, despite its extreme body mass and long extinction duration, is penalized primarily because of the combined habitat and technical challenges, whereas the thylacine benefits from lower technical difficulty and minimal risk of habitat mismatch

#### Model Deliverables

Outputs comprise (i) a ranked species list with priority bands, and (ii) sensitivity plots exploring *γ* and *θ* ranges (see Supplementary Fig. S2). All implementation code (R 4.3) and data tables are archived at Zenodo: https://doi.org/10.5281/zenodo.15924530.

## 4 Results

### 4.1 Funding Trends and Crowding-Out Assessment

Our analysis of conservation funding trends between 2021 and 2024 reveals no visual or descriptive evidence supporting the crowding-out hypothesis (Graph 1). On the contrary, the emergence of de-extinction initiatives, financed entirely through private-sector investments, coincides with an overall increase in the total funding directed toward conservation biology. All de-extinction investments during this period originated from private venture capital or philanthropic sources, whereas classical conservation and conservation biotechnology funding primarily came from government and NGO programs. Years associated with active de-extinction projects showed substantial increases in conservation-related funding without parallel declines in classical conservation budgets. Notably, 2024 saw a marked rise in conservation biotechnology funding, largely attributable to the establishment of the Colossal Foundation, which mobilized significant philanthropic investment. The proportional distribution of total funding across classical conservation, conservation biotechnology, and de-extinction is summarized in Supplementary Figure 1,2,3,4, where pie charts illustrate the percentage breakdown of total funds and the specific contributions from private, governmental, NGO, and philanthropic sources within each sector. This visualization offers a clear overview of how conservation financing is partitioned and underscores the differentiated origins of support for each conservation stream.

**Figure 1:**
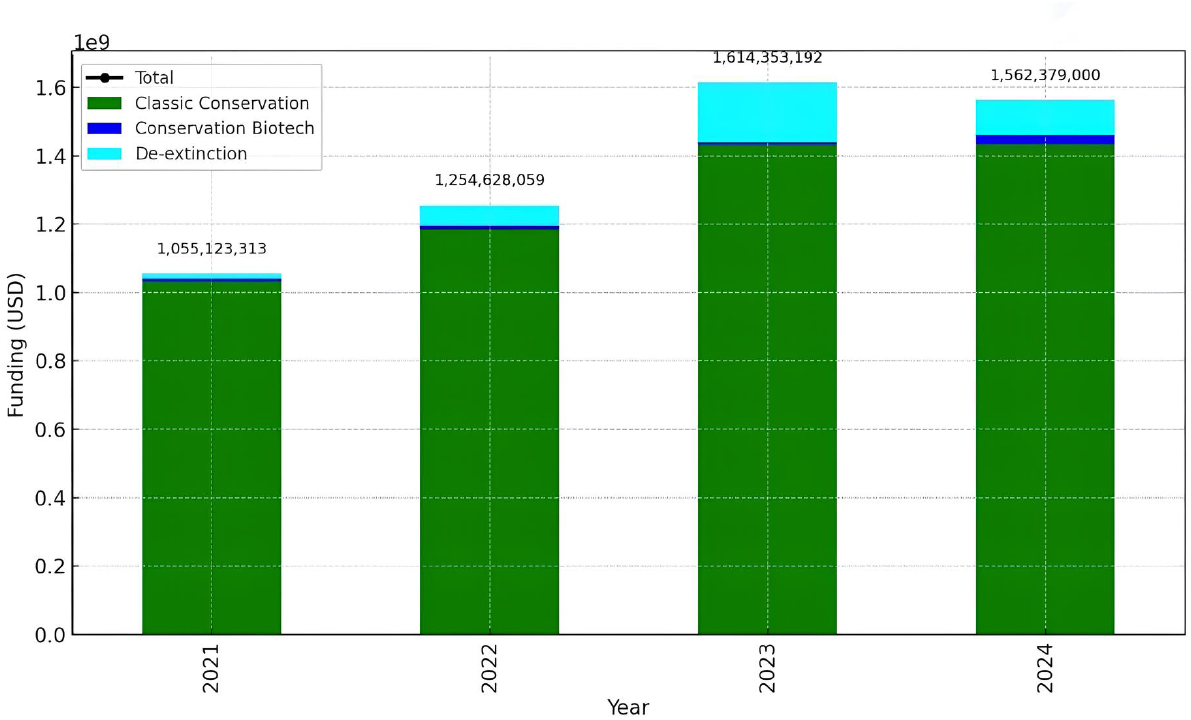
Graph-1. Annual conservation funding trends (2021–2024) by sector: classical conservation, conservation biotechnology, and de-extinction.

**Figure 2:**
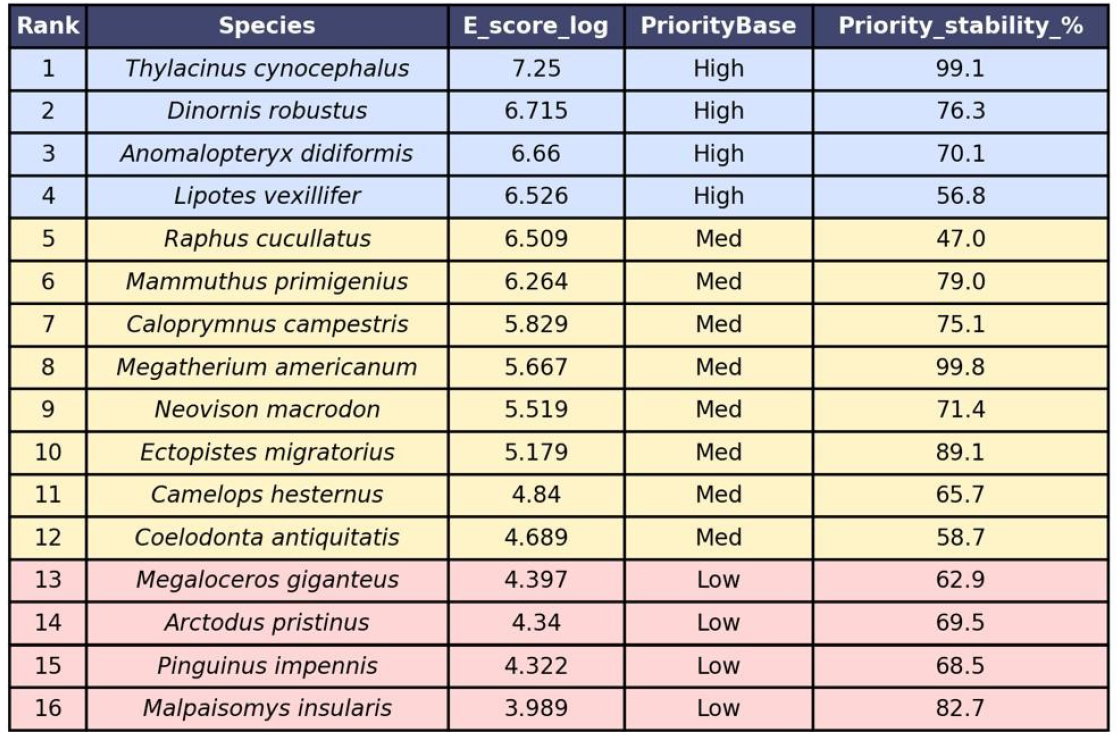
Table 1 shows that *Thylacinus cynocephalus* ranked highest, combining strong multidimensional effectiveness with exceptional stability (99.1%). Other high-priority species, including *Dinornis robustus* and *Anomalopteryx didiformis*, exhibited moderately lower stability (76.3% and 70.1%, respectively). Notably, *Lipotes vexillifer* displayed high baseline scores but reduced stability (56.8%). Within the medium-priority group, *Megatherium americanum* showed near-perfect stability (99.8%). Among low-priority taxa, *Malpaisomys insularis* (82.7%) demonstrated unexpected consistency.

**Figure 3:**
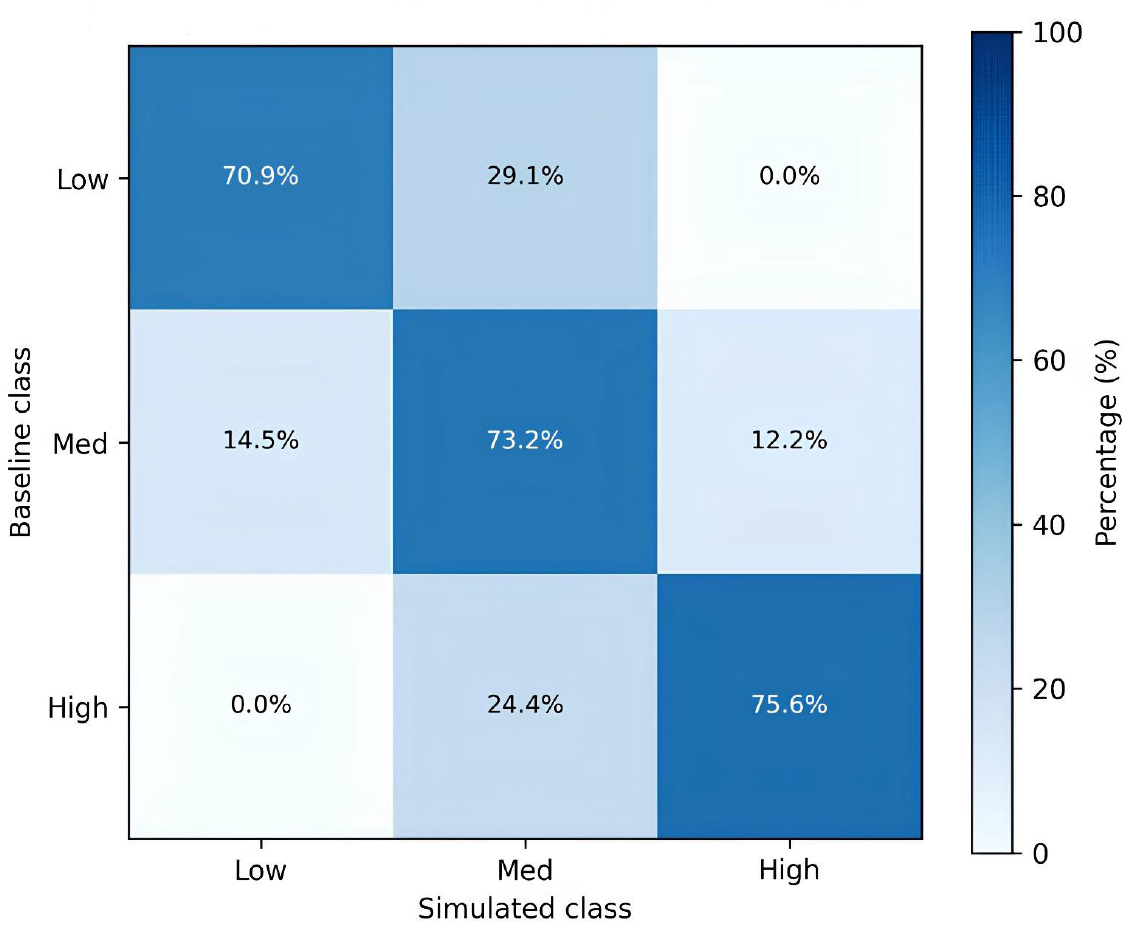
Priority-class transition heatmap (±3 jitter, 10 k runs)

### 4.2 Model outputs and overall performance

The prioritisation results showed that across all 16 species, the median class-level stability was 73.2%. *Raphus cucullatus* (dodo) showed only 47% consistency, representing the lowest overall stability in the dataset. Summary results for all taxa are provided in Figure 2.

### 4.3 Species-level results and class transitions

The priority-class transition heatmap (**Figure 3**) revealed that 70.9% of low-priority taxa remained in their class, while 29.1% occasionally shifted upward to medium; none transitioned directly to high. Medium-priority taxa were the most volatile, with 73.2% remaining medium, 14.5% dropping to low, and 12.2% rising to high. High-priority taxa were the most stable, with 75.6% retained as high and 24.4% transitioning to medium; no species moved directly from high to low.

The stability barplot (see Supplementary Figure 5) highlighted the consistently stable performance of *Megatherium americanum, Thylacinus cynocephalus*, and *Ectopistes migratorius*, while *Raphus cucullatus* and *Lipotes vexillifer* showed the lowest stability (47% and 57%, respectively). Together, these results underline the importance of integrating stability metrics into prioritisation to identify robust conservation or de-extinction candidates.

## Discussion

This study set out to examine whether the rise of de-extinction initiatives has affected the financial landscape of conservation biology. Our findings show no evidence that de-extinction projects are taking resources away from traditional conservation. Instead, they contribute additional funding, mainly from private investors, thereby increasing the total funding available for conservation. This challenges the common claim that money spent on de-extinction would be better used for protecting existing species. Our data show that the money going into deextinction comes from completely different sources. It is largely funded by private individuals and organizations interested in technological breakthroughs, who would likely not invest in classical conservation projects. In other words, these are new funds that would not have gone to conventional conservation anyway. Redirecting these funds to traditional conservation is not just unlikely; it also overlooks that these investors are motivated by technological innovation, and their support expands rather than replaces existing conservation efforts.

While our results suggest that de-extinction projects currently do not take resources away from traditional conservation, this does not guarantee that future developments will have no financial impact. Moreover, it remains unclear what level of funding would be required for the reintroduction, management, and long-term monitoring of resurrected species in the wild. These downstream costs, which could include habitat restoration, ongoing veterinary care, and potential conflict mitigation, are difficult to estimate at present and represent an important area for future research.

The basic idea of de-extinction is to bring back extinct species, but with today’s technologies, it is still not possible to fully recreate an organism from scratch. Rather than aiming for exact genetic replication, the primary objective of de-extinction science is to produce ecologically functional equivalents capable of fulfilling comparable ecological roles [3]. In this context, the announcement in 2025 by Colossal Biosciences of the creation of proxy dire wolves sparked significant scientific debate [24]. Although these animals were presented as “dire wolves,” they are in fact dire wolf-like organisms, fitting the definition of de-extinction as they aim to recreate functional equivalents, not exact genetic copies. Nonetheless, while some researchers refer to them directly as dire wolves, others describe them merely as genetically modified wolves [21, 23]. This disagreement has not only highlighted a broader classification challenge in the field, namely how to categorize organisms that are synthetically produced or re-created through de-extinction technologies, but has also raised another important issue regarding how society and science define the identity and authenticity of such organisms. These debates tie into broader concerns about the societal and ethical implications of de-extinction. Because of this, it is crucial to avoid framing de-extinction as a justification for tolerating further extinctions; using the idea of resurrection as an excuse to allow more species loss would be a serious ecological and ethical mistake. As argued by [25], extinct species have symbolic value as martyrs for the conservation cause, and de-extinction risks eroding the moral urgency associated with extinction events. De-extinction should never be seen as a safety net that makes species loss acceptable or reduces the urgency of protecting threatened species. Managing this risk of false reassurance is essential.

The proposed 2025 budget cuts to conservation biology in the United States may already signal the start of such dangerous complacency [20]. Contrary to claims that present de-extinction as a rival to traditional conservation, the scale and urgency of today’s biodiversity crisis call for integration, not division. De-extinction should be seen as an innovative branch of conservation biology that adds to, rather than replaces, other efforts. The future of biodiversity protection depends on finding the right balance and ensuring that new technologies complement, rather than undermine, classical conservation strategies.

At the same time, as in traditional conservation programs, there is a risk that de-extinction efforts will disproportionately focus on charismatic species, driven by their ability to attract funding and public attention. If such projects are pursued solely as commercial ventures or zoo attractions, they should not be framed as conservation efforts. While it is well established that charismatic species can help mobilize public interest, recent work [22] highlights that these species are often selected not just for their symbolic appeal but through structural biases shaped by the interactions between researchers, funders, and user groups. This raises the risk that conservation priorities are skewed away from ecological or restoration relevance, reinforcing patterns of charisma-driven bias. The challenge, therefore, is to balance scientific and ecological priorities with public engagement, ensuring that conservation strategies are guided by evidence-based and ethical principles, not merely by marketability or fundraising potential. Ultimately, the challenge is not only to allocate funds more accurately but also to enlarge the overall conservation budget. Even so, mathematical models remain indispensable for weighing priorities in a transparent and repeatable way. The framework presented here ranks de-extinction candidates by integrating ecological value, technological upside, and symbolic or strategic impact. Our results show that projects already benefitting from active research, development, and funding, such as the thylacine, woolly mammoth, and moa, emerge as the most rational and highest-scoring options within a necessarily limited candidate pool. However, species such as the dodo warrant more careful consideration, given both the relative instability of their ranking in our model and the concerns raised in recent literature [22] about charisma-driven selection biases in de-extinction efforts.

This underscores that while quantitative prioritization tools provide valuable guidance, they must be situated within a broader ethical and ecological framework to ensure that decision-making reflects both scientific evidence and societal values. The model does not attempt to answer the broader ethical question of whether de-extinction should be pursued. Instead, using current economic data, it shows two key points. First, de-extinction initiatives bring new resources into conservation biology. Second, the species currently under development are, based on our metrics, genuinely high-value targets. With better parameter tuning and the addition of species-specific conservation data, future versions of the model could help optimize both de-extinction projects and priorities for living species, reducing waste in a field where grants and donations are limited. More broadly, similar models could also be adapted for use in classical conservation programs, helping guide decisions on how to use resources more effectively across conservation efforts.

Mathematical models can never mirror the real world perfectly; their variables are abstractions by definition. Public support will likely grow as the everyday consequences of biodiversity loss become more visible, yet the goal is to curb that loss before the impacts intensify. Our findings suggest that conservation biotechnology, including its de-extinction branch, has considerable potential to attract funding, but this potential should complement traditional conservation finance rather than cannibalise it.

As noted in the Methods section, the habitat degradation scores (*d*_*i*_) used in our model are intentionally coarse, designed as heuristic estimates to capture broad patterns of ecosystem transformation. A clear avenue for improvement would be to replace these author-assigned scores with quantitative metrics derived directly from geospatial data layers specific to the candidate species’ historical and potential reintroduction areas. High-resolution land-cover change maps, climate data, species distribution models, and human footprint indices could be integrated to generate a composite habitat degradation index. In addition, most variables in the current framework, including biodiversity and symbolic value scores, could be refined using richer data sources and field studies, such as targeted habitat assessments to evaluate the suitability of reintroduction sites. Importantly, similar modeling approaches could also be applied beyond de-extinction, helping to improve decision-making and resource allocation in classical conservation programs as well.

Building on the insights of [12] and [23], we suggest that de-extinction should be understood as an advanced but expensive instrument, to be applied only in those rare and complicated situations where the ecological benefits are clear, the ethical case is sound, and long-term governance structures are in place; its ultimate worth will be measured not by the spectacle of resurrecting charismatic species, but by the degree to which it strengthens public engagement, catalyses technological innovation, and, most importantly, supports rather than supplants the foundational practices of conservation biology that remain essential for safeguarding Earth’s living diversity.

## Supporting information

Supplementary Figures and Supplementary Material: Data Sources (PDF)

