## Supplementary Figures and Supplementary Material: Data Sources (PDF) for "Rethinking De-Extinction Criticism: A Multi-Dimensional Model for Prioritizing Revivable Species under Funding Controversies"

Alper Kaan Selçukoğlu


Deha Kaykı


July 16, 2025

### Data Sources and References

- Full datasets and code: Zenodo archive (https://doi.org/10.5281/zenodo.15924530)


- Allentoft, M. E., Heller, R., Holdaway, R. N., & Bunce, M. (2015). Ancient DNA microsatellite analyses of the extinct New Zealand giant moa (*Dinornis robustus*) identify relatives within a single fossil site. *Heredity*, 115(6), 481–487. https://doi.org/10.1038/hdy.2015.48

- Hughes, S., Hayden, T. J., Douady, C. J., Tougard, C., Germonpré, M., Stuart, A., Lbova, L., Carden, R. F., Hänni, C., & Say, L. (2006). Molecular phylogeny of the extinct giant deer, *Megaloceros giganteus*. *Molecular Phylogenetics and Evolution*, 40(1), 285–291. https://doi.org/10.1016/j.ympev.2006.02.004

- Firmat, C., Rodrigues, H. G., Hutterer, R., Rando, J. C., Alcover, J. A., & Michaux, J. (2010). Diet of the extinct Lava mouse *Malpaisomys insularis* from the Canary Islands: insights from dental microwear. *The Science of Nature*, 98(1), 33–37. https://doi.org/10.1007/s00114-010-0738-z

- Sealfon, R. A. (2007). Dental divergence supports species status of the extinct sea mink (*Carnivora: Mustelidae: Neovison macrodon*). *Journal of Mammalogy*, 88(2), 371–383. https://doi.org/10.1644/06-mmm-a-227r1.1

- Cai, Z., Petersen, B., Sahana, G., Madsen, L. B., Larsen, K., Thomsen, B., Bendixen, C., Lund, M. S., Guldbrandtsen, B., & Panitz, F. (2017). The first draft reference genome of the American mink (*Neovison vison*). *Scientific Reports*, 7(1). https://doi.org/10.1038/s41598-017-15169-z

- Wang, L., Zhou, S., Lyu, T., Shi, L., Dong, Y., He, S., & Zhang, H. (2022). Comparative genome analysis reveals the genomic basis of semiaquatic adaptation in American mink (*Neovison vison*). *Animals*, 12(18), 2385. https://doi.org/10.3390/ani12182385

- Heintzman, P. D., Zazula, G. D., Cahill, J. A., Reyes, A. V., MacPhee, R. D., & Shapiro, B. (2015). Genomic data from extinct North American *Camelops* revise camel evolutionary history. *Molecular Biology and Evolution*, 32(9), 2433–2440. https://doi.org/10.1093/molbev/msv128

- Mitchell, K. J., Bray, S. C., Bover, P., Soibelzon, L., Schubert, B. W., Prevosti, F., Prieto, A., Martin, F., Austin, J. J., & Cooper, A. (2016). Ancient mitochondrial DNA reveals convergent evolution of giant short-faced bears (*Tremarctinae*) in North and South America. *Biology Letters*, 12(4), 20160062. https://doi.org/10.1098/rsbl.2016.0062

- Rovinsky, D. S., Evans, A. R., & Adams, J. W. (2019). The pre-Pleistocene fossil thylacinids (Dasyuromorphia: Thylacinidae) and the evolutionary context of the modern thylacine. *PeerJ*, 7, e7457. https://doi.org/10.7717/peerj.7457

- Yuan, J., Sheng, G., Hou, X., et al. (2014). Ancient DNA sequences from *Coelodonta antiquitatis* in China reveal its divergence and phylogeny. *Science China Earth Sciences*, 57, 388–396. https://doi.org/10.1007/s11430-013-4702-6

- Pagès, M., Chevret, P., Gros-Balthazard, M., Hughes, S., Alcover, J. A., Hutterer, R., Rando, J. C., Michaux, J., & Hänni, C. (2012). Paleogenetic analyses reveal unsuspected phylogenetic affinities between mice and the extinct *Malpaisomys insularis*, an endemic rodent of the Canaries. *PLoS ONE*, 7(2), e31123. https://doi.org/10.1371/journal.pone.0031123

- Thomas, J. (2018). Evolution & Extinction of the Great Auk: A Palaeogenomic Approach. Student thesis: Doctor of Philosophy.

- Shapiro, B., Sibthorpe, D., Rambaut, A., Austin, J., Wragg, G. M., Bininda-Emonds, O. R. P., Lee, P. L. M., & Cooper, A. (2002). Flight of the Dodo. *Science*, 295(5560), 1683. https://doi.org/10.1126/science.295.5560.1683

- Westerman, M., Loke, S., & Springer. (2003). Molecular phylogenetic relationships of two extinct potoroid marsupials, *Potorous platyops* and *Caloprymnus campestris* (Potoroinae: Marsupialia). *Molecular Phylogenetics and Evolution*, 31(2), 476–485. https://doi.org/10.1016/j.ympev.2003.08.006

- Edwards, S. V., Cloutier, A., Cockburn, G., Driver, R., Grayson, P., Katoh, K., Baldwin, M. W., Sackton, T. B., & Baker, A. J. (2024). A nuclear genome assembly of an extinct flightless bird, the little bush moa. *Science Advances*, 10(21). https://doi.org/10.1126/sciadv.adj6823

- Drucker, D. G. (2022). The isotopic ecology of the mammoth steppe. *Annual Review of Earth and Planetary Sciences*, 50(1), 395–418. https://doi.org/10.1146/annurev-earth-100821-081832

- Miller, W., Drautz, D. I., Ratan, A., Pusey, B., Qi, J., Lesk, A. M., Tomsho, L. P., Packard, M. D., Zhao, F., Sher, A., Tikhonov, A., Raney, B., Patterson, N., Lindblad-Toh, K., Lander, E. S., Knight, J. R., Irzyk, G. P., Fredrikson, K. M., Harkins, T. T., . . . Schuster, S. C. (2008). Sequencing the nuclear genome of the extinct woolly mammoth. *Nature*, 456(7220), 387–390. https://doi.org/10.1038/nature07446

- Kuzmin, Y. V. (2009). Extinction of the woolly mammoth (*Mammuthus primigenius*) and woolly rhinoceros (*Coelodonta antiquitatis*) in Eurasia: Review of chronological and environmental issues. *Boreas*, 39(2), 247–261. https://doi.org/10.1111/j.1502-3885.2009.00122.x

- Orlando, L., Leonard, J. A., Thenot, A., Laudet, V., Guerin, C., & Hänni, C. (2003). Ancient DNA analysis reveals woolly rhino evolutionary relationships. *Molecular Phylogenetics and Evolution*, 28(3), 485–499. https://doi.org/10.1016/s1055-7903(03)00023-x

- Figueirido, B., & Janis, C. M. (2011). The predatory behaviour of the thylacine: Tasmanian tiger or marsupial wolf? *Biology Letters*, 7(6), 937–940. https://doi.org/10.1098/rsbl.2011.0364

- Lord, E., Dussex, N., Kierczak, M., Díez-Del-Molino, D., Ryder, O. A., Stanton, D. W., Gilbert, M. T. P., Sánchez-Barreiro, F., Zhang, G., Sinding, M. S., Lorenzen, E. D., Willerslev, E., Protopopov, A., Shidlovskiy, F., Fedorov, S., Bocherens, H., Nathan, S. K., Goossens, B., Van Der Plicht, J., . . . Dalén, L. (2020). Pre-extinction demographic stability and genomic signatures of adaptation in the woolly rhinoceros. *Current Biology*, 30(19), 3871–3879.e7. https://doi.org/10.1016/j.cub.2020.07.046

- Feigin, C., Frankenberg, S., & Pask, A. (2022). A chromosome-scale hybrid genome assembly of the extinct Tasmanian tiger (*Thylacinus cynocephalus*). *Genome Biology and Evolution*, 14(4). https://doi.org/10.1093/gbe/evac048

- Delsuc, F., Kuch, M., Gibb, G. C., Karpinski, E., Hackenberger, D., Szpak, P., Martínez, J. G., Mead, J. I., McDonald, H. G., MacPhee, R. D., Billet, G., Hautier, L., & Poinar, H. N. (2019). Ancient mitogenomes reveal the evolutionary history and biogeography of sloths. *Current Biology*, 29(12), 2031–2042.e6. https://doi.org/10.1016/j.cub.2019.05.043

- Zhang, X., Wang, D., Liu, R., Wei, Z., Hua, Y., Wang, Y., Chen, Z., & Wang, L. (2003). The Yangtze River dolphin or baiji (*Lipotes vexillifer*): Population status and conservation issues in the Yangtze River, China. *Aquatic Conservation: Marine and Freshwater Ecosystems*, 13(1), 51–64. https://doi.org/10.1002/aqc.547

- Zhou, X., Sun, F., Xu, S., Fan, G., Zhu, K., Liu, X., Chen, Y., Shi, C., Yang, Y., Huang, Z., Chen, J., Hou, H., Guo, X., Chen, W., Chen, Y., Wang, X., Lv, T., Yang, D., Zhou, J., . . . Yang, G. (2013). Baiji genomes reveal low genetic variability and new insights into secondary aquatic adaptations. *Nature Communications*, 4(1). https://doi.org/10.1038/ncomms3708

- Murray, G. G. R., Soares, A. E. R., Novak, B. J., Schaefer, N. K., Cahill, J. A., Baker, A. J., Demboski, J. R., Doll, A., Da Fonseca, R. R., Fulton, T. L., Gilbert, M. T. P., Heintzman, P. D., Letts, B., McIntosh, G., O'Connell, B. L., Peck, M., Pipes, M., Rice, E. S., Santos, K. M., ... Shapiro, B. (2017). Natural selection shaped the rise and fall of passenger pigeon genomic diversity. *Science*, 358(6365), 951–954. https://doi.org/10.1126/science.aao0960

- Ellsworth, J. W., & McComb, B. C. (2003). Potential effects of passenger pigeon flocks on the structure and composition of presettlement forests of eastern North America. *Conservation Biology*, 17(6), 1548–1558. https://doi.org/10.1111/j.1523-1739.2003.00230.x

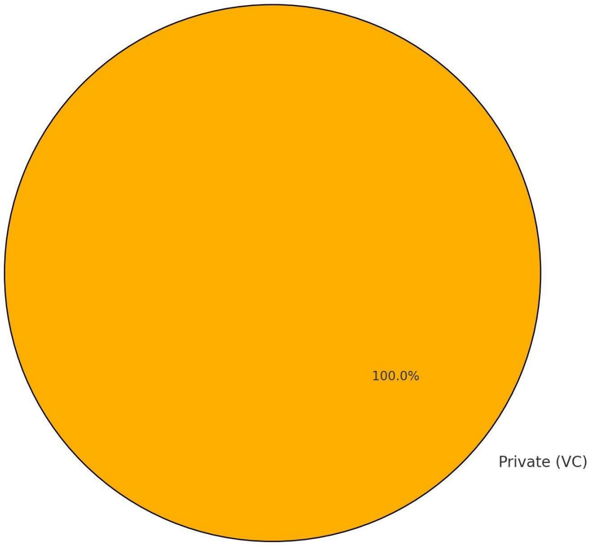


Figure 1: Funding type distribution for de-extinction projects, showing that 100% of funding originates from private venture capital (VC) sources.

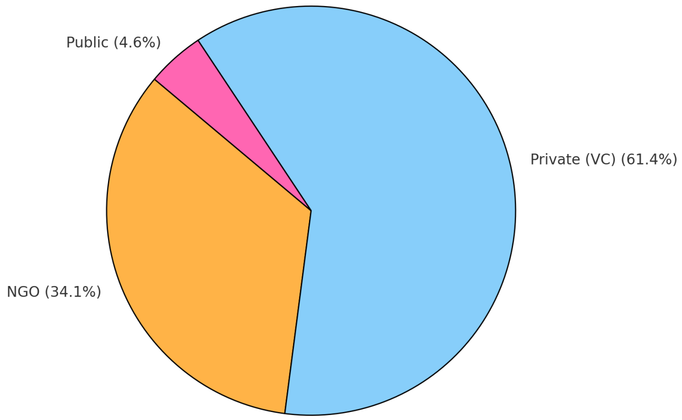


Figure 2: Funding type distribution for Conservation Biotech based on total financial contributions, showing that 61.4% of the funding originated from Private (VC) sources, 34.1% from NGOs, and 4.6% from Public sources.

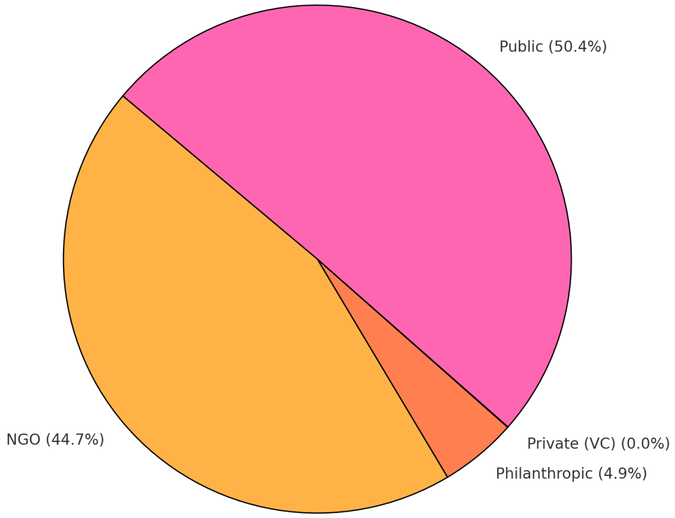


Figure 3: Funding type distribution for Classic Conservation based on total financial contributions, showing that 50.4% of the funding originated from Public sources, 44.7% from NGOs, 4.9% from Philanthropic sources, and 0.0% from Private (VC) sources.

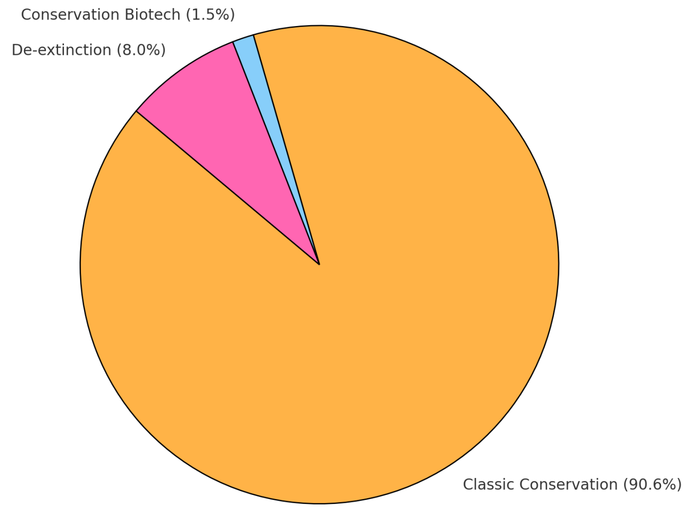


Figure 4: Overall funding distribution by field, showing that 90.6% of the total funding was allocated to Classic Conservation, 8.0% to De-extinction, and 1.5% to Conservation Biotech.

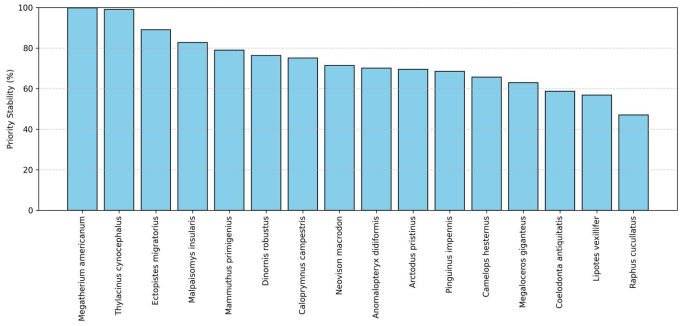


Figure 5: Monte-Carlo simulation results showing priority class stability under random perturbation (±3 jitter, 10k runs). Species are ranked by their stability percentage, highlighting robustness across priority classifications.
